# *Pontoscolex corethrurus:* a Homeless Invasive Tropical Earthworm?

**DOI:** 10.1101/624122

**Authors:** Angel I. Ortíz-Ceballos, Diana Ortiz-Gamino, Antonio Andrade-Torres, Paulino Pérez-Rodríguez, Maurilio López-Ortega

## Abstract

The presence of earthworm species in crop fields is as old as agriculture itself. The earthworms *Pontoscolex corethrurus* (invasive) and *Balanteodrilus pearsei* (native) are associated with the emergence of agriculture and sedentism in the region Amazon and Maya, respectively. Both species have shifted their preference from their natural habitat to the cropland niche; however, they contrast in terms of intensification of agricultural land use (anthropic impact to the symbiotic soil microbiome). *P. corethrurus* inhabits conventional agroecosystems (pesticides, herbicides, and fertilizers are applied to soil), while *B. pearsei* thrives in traditional agroecosystems (biological management of soil); that is, *P. corethrurus* has not yet been recorded in soils where *B. pearsei* dwells. The demographic behavior of these two earthworm species was assessed in the laboratory over 100 days, according to their origin (OE; *P. corethrurus* and *B. pearsei*) food quality (FQ; soil only, maize stubble, and *Mucuna pruriens*), and soil moisture (SM; 25, 33, and 42%). The results showed that OE, FQ, SM, and the OE x FQ interaction were highly significant for the survival, growth, and reproduction of earthworms. *P. corethrurus* showed a lower survival rate (> mortality). *P. corethrurus* survivors fed a diet of low-to-intermediate nutritional quality (soil and corn stalks, respectively) showed a greater capacity to grow and reproduce; however, it was surpassed by the native earthworm when fed a high-quality diet (*M. pruriens*). Besides, *P. corethrurus* displayed a low cocoon hatching (emergence of juveniles). These results suggest that the presence of the invasive species was associated with the absence of natural mutualistic bacteria (gut, nephridia, and cocoons), and with a negative interaction with the soil microbiota where the native species dwells. These results are consistent with the absence of *P. corethrurus* in milpa and pasture-type agricultural niches managed by peasants (agroecologists) to grow food regularly a biological management of soil. The results reported here and the published information jointly (e.g., designation of the neotype and ambiguity of the place of origin) jointly suggest that *P. corethrurus* is an invasive species that is neither wild nor domesticated, that is, its eco-evolutionary phylogeny needs to be derived based on its symbionts.

## Introduction

Although humans have produced novel niches prior to the advent of agriculture, the innovation of domestication led to changes in the life cycle of one or a few species, and the local microenvironments were manipulated, especially soil biota (Bender et al. 2016; Boivin et al. 2016; Pérez-Jaramillo et al. 2016; Fuller and Ducas 2017). The artificial landscapes that resulted from these practices (anthropocentric ecology) were exported as agricultural packages from the centers of origin (Bender et al 2016; Boivin et al. 2016; Fuller and Ducas 2017). Thus, over a relatively short period in the history of mankind, the expansion of agriculture has brought about the remodeling of biodiversity as one of the most significant anthropogenic impacts on terrestrial ecosystems (Lodge 1993; Boivin et al. 2016; Bender et al. 2016).

Agriculture has given rise to uniform and predictable disturbed ecological niches (invasible habitats), which have proven highly beneficial for non-domesticated species or weeds (Boivin et al. 2016; Bender et al. 2016; Fuller and Stevens 2017), and some earthworm species. Blakemore (2009) has suggested that the origins of cosmopolitan (invasive) earthworms at family level are associated with domestication centers of plants and animals; that is, the presence of earthworms in crop fields is as old as agriculture itself (Blakemore 2009; Vigueira et al. 2013; Mercuri et al. 2018). The terms of the Millennium Ecosystem Assessment highlight the catalytic role of earthworms regarding two environmental services (Plaas et al. 2019), namely the formation of soil and biogeochemical cycles, both of which are prerequisites for other environmental services (Brussaard et al. 2012; Plaas et al. 2019).

Most of the studies focused on earthworms have used species adapted to crops, and most of them are currently considered as invasive (Brussaard et al. 2012). It has been documented that 3 % (100-120) of the diversity of earthworms are invasive species (Hendrix 2008), and stating that just a few of them had a negative impact on terrestrial agroecosystems would not be an exaggeration (Simberloff 2009; Hendrix et al. 2006). As an example, European earthworms are frequently mentioned as the main cause of an irreversible change in the diversity and functioning of ecosystems in North America (Wisconsin glaciation areas) that were previously free from earthworms, 12 thousand years ago (Uvarov 2009; Lobe et al. 2014; Vastergard et al. 2015). However, there is a deeply rooted positive attitude toward earthworms in human populations in North America, acknowledging their beneficial effects on agricultural soils and urban gardens (Simmons et al. 2015; Plaas et al. 2019).

Among the invasive tropical earthworms, the endogeic species *Pontoscolex corethrurus* was collected and described in crop fields in Blumenau, Brazil 160 years ago (Müller 1857; James et al. 2019); it has a broad distribution range and is the most studied tropical species (Fragoso 2018; Taheri et al. 2018a). Native species also move across a region in a similar way to invasive species, in addition to natural displacements (Blakemore 2009; Nackley et al. 2017). The native endogeic earthworm *Balanteodrilus pearsei* was first collected and described from Gongora cave in Okcutzcab, Yucatan 81 years ago (Orell 2019); it is distributed in the east and southeast of Mexico and Belize (Fragoso 2018); it dwells in natural and agricultural environments and is the most studied species native to Mexico. Most studies conducted with both species point to a positive influence of their biological activity on soil (Taheri et al. 2018b), i.e., they do not meet the definition of pest (Richardson et al. 2000). For this reason, we use the term *invasive* with reference to the biogeographical status of the species, regardless of its impact on soil (Richardson et al. 2000; Ricciardi and Cohen 2007).

Similar to weeds (Willcox 2012; Vigueira et al. 2013; Fuller and Stevens 2017; Mercuri et al. 2018), it can be suggested that *P. corethrurus* and *B. pearsei* have shifted their preference from their natural habitat to agricultural environments, spreading geographically beyond their place of origin, and are currently key elements of agricultural environments. The presence of *P. corethrurus* and *B. pearsei* is associated with the development of pre-Columbian cultivation techniques in the Amazon (Clement et al. 2015; Levis et al. 2018; Boivin et al. 2016; Watling et al. 2018) and Maya (Ford and Nigh 2009; McNeil 2012; Boivin et al. 2016) regions, respectively; for example, it is believed that *P. corethrurus* facilitated the formation of fertile soils in the Amazon area named “Terra Preta do Indo” (Glaser et al. 2000; Ponge 2006). Both species have adapted to niches that emerged from agriculture (Laland et al. 199), but contrast regarding the intensification of agricultural land use and/or the diversion of each from natural habitats (anthropic manipulation of soil). *P. corethrurus* is commonly found in conventional agrecosystems (use of fertilizers, herbicides, pesticides, and tillage), as well as in industrial (polluted with heavy metals, petroleum hydrocarbons, and others) and urban areas (Lavelle et al. 1987; Fragoso et al. 2018; Marichal et al. 2010; Taheri et al. 2018a). For its part, *B. pearsei* inhabits soils managed under an agroecological approach (little human impact of the soil microbiome), such as traditional agroecosystems (no use of industrial inputs) and in natural ecosystems (; Ortiz-Ceballos et al. 2004; Huerta et al. 2006; Fragoso et al. 2016). *P. corethrurus* has been found coexisting with native species in some agroecosystems (Marichal et al. 2010; Ortiz-Gamino et al. 2016), but there are no records of its coexistence with *B. pearsei* so far (Ortiz-Ceballos et al. 2004; Huerta et al. 2006; Fragoso et al. 2016).

A previous study of coexistence under controlled conditions showed no competitive interaction between *P. corethrurus* and *B. pearsei*, i.e., both can coexist (Ortiz-Ceballos et al. 2005). However, the question to address is, why *P. corethrurus* has not invaded the agroecological niche of *B. pearsei*? Therefore, this work compared the demographic behavior of *P. corethrurus* vs. *B. pearsei* assuming that the survival rate of the invasive species decreases in soil populated by the native species.

## Materials and Methods

The experimental procedure used in this study is detailed elsewhere (Ortiz-Ceballos et al., 2005).

### Soil

Soil was collected from a maize field (MM) rotated with the tropical legume velvet bean (*Mucuna prurien var. utilis*). The silty clay loam soil (41.5 % sand; 26.8 % clay; 31.6 % silt) was air-dried in the shade at room temperature and sieved through a 2 mm mesh. The main chemical characteristics of this soil were: 2.7 % organic matter; 0.14 % total N; 11.4 C/N; pH (H_2_O) of 6.3.

### Earthworms

Two tropical endogeic earthworm species were used in this study: *B. pearsei* (native) and *P. corethrurus* (invasive). *B. pearsei* was collected from the MM field, whereas *P. corethrurus* was collected from pastures, given its absence in the former site. All earthworms (120 for each species) were collected two weeks prior to the beginning of the experiment.

### Food Quality

The effects of foof quality were assessed by using two different types of plant litter of contrasting nutritional quality: *M. pruriens* (52.4 % C, 2.25 % N, 23.3 C/N, and 9.67 % ash) and maize (52 % C, 0.84 % N, 61.9 % C/N, and 10.3 % ash). Both materials were obtained from the MM field, oven-dried at 60 °C for 48 h, and sieved (1 mm).

### Experiment

Growth, sexual maturity, reproduction (cocoons and juveniles), and mortality of *B. pearsei* and *P. corethrurus* were investigated during 100 days using a factorial design with three factors: origin of earthworms (OE), soil moisture (SM), and food quality (FQ). SM involved 3 levels, corresponding to the permanent wilt point (25 %), field capacity (42 %), and an intermediate level (33 %). FQ included three levels: 300 g soil only (S), 294 g soil + 6 g maize stubble (MS), and 294 g soil + 6g *M. pruriens* (MP); the amounts added correspond to those commonly found in both maize monocultures and cultures rotating maize and *M. pruriens.* The earthworm species used belong to two different classes based on origin: Native (*B. pearsei*) and Invasive (*P. corethrurus*).

The combination of the three factors and three levels produced nine treatments with five replicates per treatment. Each replicate consisted of a plastic container (12×12×8 cm) containing 300 g dried soil of the corresponding food-soil mixture and soil moisture; two individuals of *B. pearsei* and two of *P. corethrurus* were transferred to each container (Table 1).

**Table 1.**
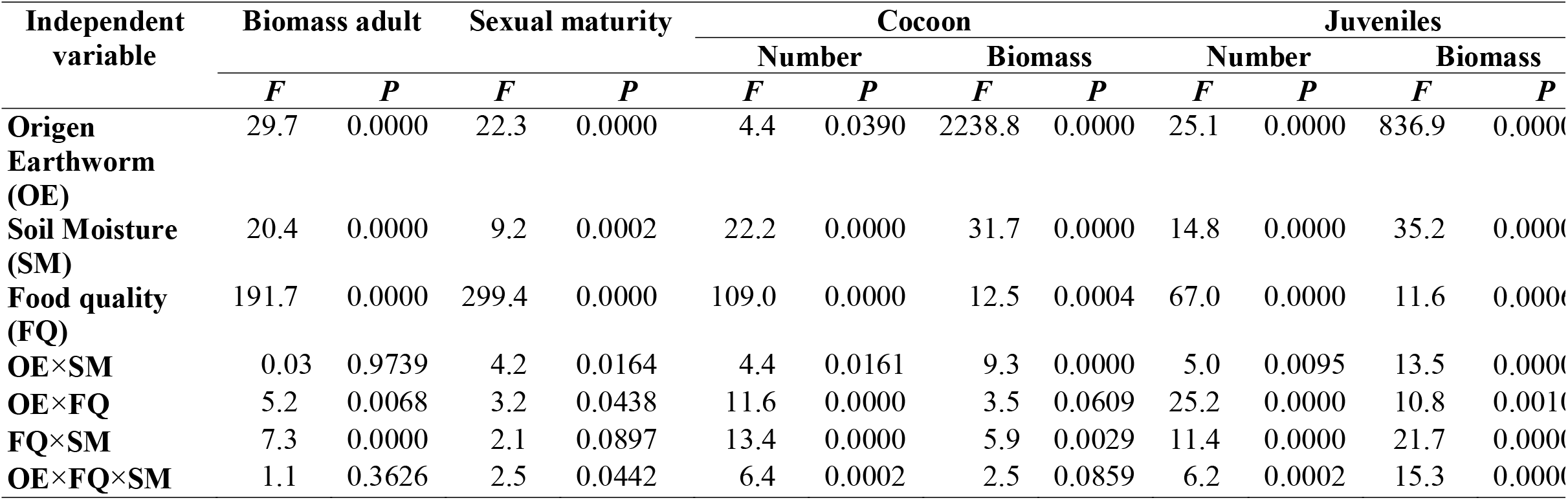
*F*-values and significance levels (ANOVA) of the interaction of three factors on growth and reproduction of t tropical endogeic earthworm *Pontoscolex corethrurus* and *Balanteodrilus pearsei* after 100 days of culture in soil w low anthropic impact

Earthworms were washed, dried on paper towels, weighed, and assigned randomly to each treatment. The baseline weight of the 45 replicates from the nine treatments was statistically similar in *B. pearsei* and *P. corethrurus* (76.06 ± 26.1 mg, n = 90 and 66.04 ± 31.1 mg, n = 90, respectively). Containers were incubated at 26 ± 1 °C. Body weight, mortality, clitellum appearance (sexual maturity), and number and biomass of cocoons and juveniles of *B. pearsei* and *P. corethrurus* were recorded at 10-day intervals, and soil was replaced. Before use, fresh soil (including the corresponding food-soil mixture and moisture) was preincubated for 8 days at 26 °C in order to trigger litter substrate decomposition. Each cocoon produced was incubated in a petri dish at 26 °C; incubation time as well as number and weight of all juveniles hatched were recorded.

### Statistical Analysis

Cocoon and juvenile weight, and growth were evaluated through Analysis of Varienace (ANOVA. Mortality, sexual maturity, number of cocoons, and number of juveniles were analyzed using generalized linear models, specifically the Poisson distribution wich is widely used for modelling count data. Differences between means were evaluated with Tukey’s HSD. All statistical analyses were perfomed using the Statistica software.

## Results

### Mortality

At 100 days of culture, significant effects were observed between the origin of earthworms (OE), food quality (FQ), soil moisture (SM) and mortality, and the interaction between these three factors on sexual maturity, number of cocoons, and number and biomass of juveniles. The invasive earthworm (*P. corethrurus*) had a 21.1 % mortality rate in the soil treatment (33 % and 25 % SM). In contrast, the native earthworm (*B. pearsei*) had only a 1.1 % mortality rate in *M. pruriens* (only 42 % SM).

### Growth

Growth of the endogeic earthworms clearly varied in response to EO, FQ, SM, and the EO×CF and SM ×CF interactions (Table 1). At 100 days of culture, the growth of the invasive and native species (*P. corethrurus* and *B. pearsei*, respectively) was higher when food quality increased (Fig. 1). In the three FQ levels (soil, maize stubble, and *M. pruriens*) the exotic species showed a faster growth (1.6, 9.4, and 12.3 mg/day, respectively) relative to the native species (0.34, 4.8, and 10.4 mg/day).

**Fig. 1.**
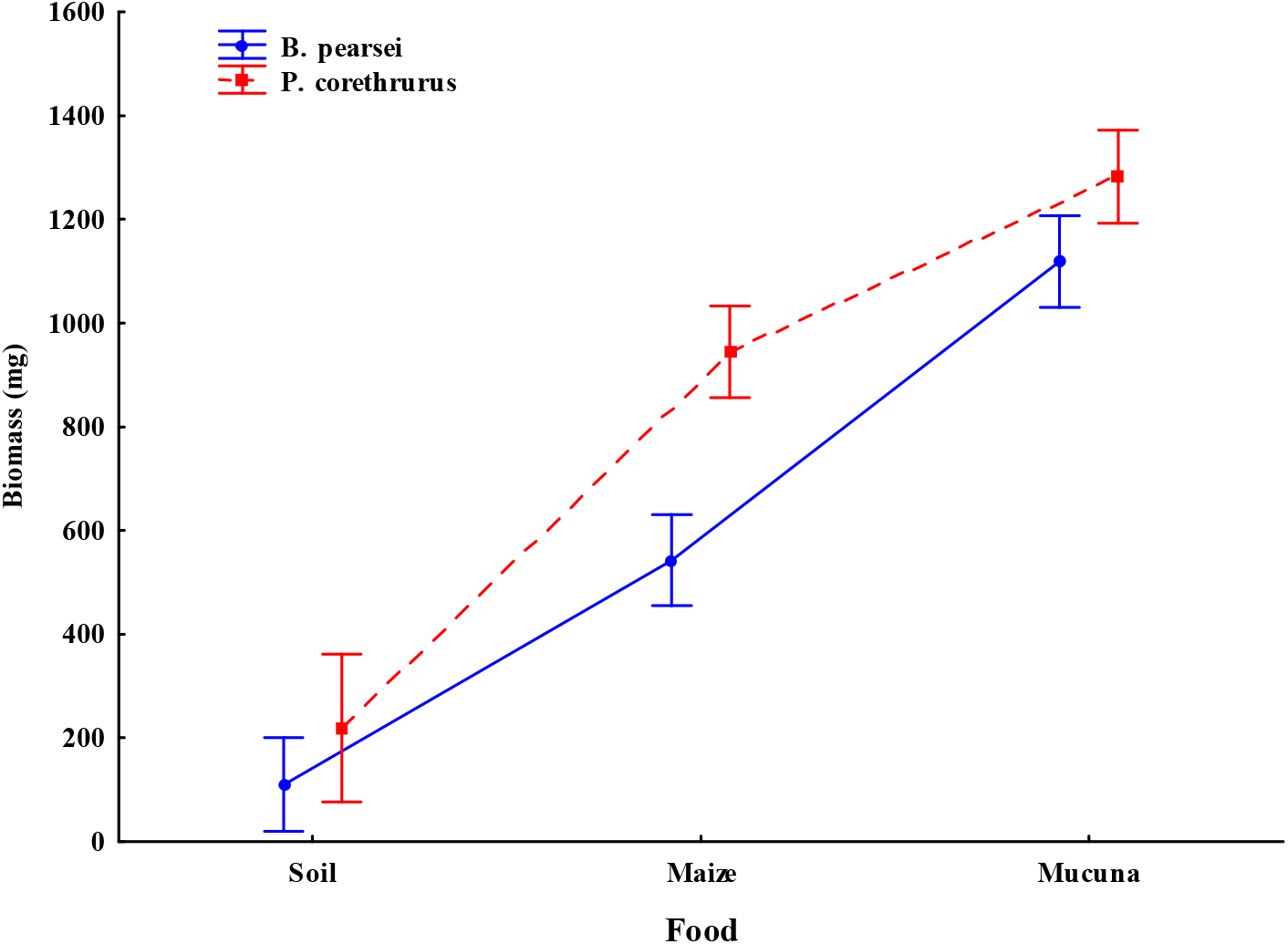
Growth rate of the tropical endogeic earthworms *Pontoscolex corethrurus* (invasive) and *Balanteodrilus pearsei* (native) after 100 days of culture using three diets of different nutritional quality in soil with low anthropic impact. Vertical lines represent standard error.

### Reproduction

#### *Sexual Maturity* (clitellum)

When fed *M. pruriens*, the onset of sexual maturity in *P. corethrurus* and *B. pearsei* occurred at 30 days; when fed maize stubble, it occurred at 30 and 70 days in *P. corethrurus* and *B. pearsei*, respectively.

At 100 days of culture, OE, FQ, SM, and the OE × CF × SM interaction significantly affected clitellum development (Table 1). The invasive and native earthworms reached sexual maturity in the treatments with *M. pruriens* (100 % and 86.6 %) and maize stubble (96.7 % and 70.0 %), respectively (Fig. 2). No individuals reached sexual maturity after 100 days in the soil treatments; however, in the soil treatment with 33 % SM, one earthworm of *P. corethrurus* (6.7 %) reached sexual maturity at 80 days.

**Fig. 2.**
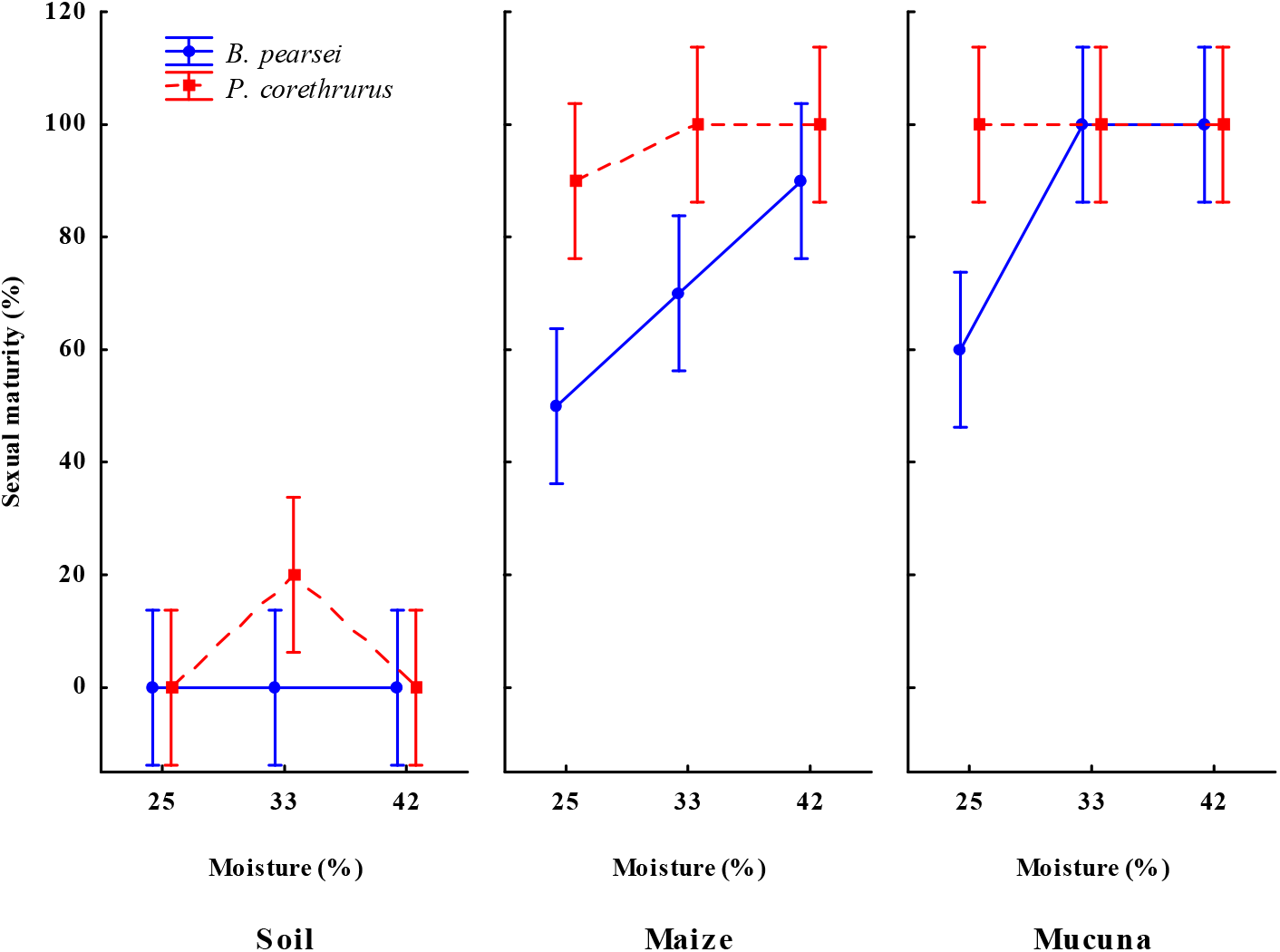
Sexual maturity (formation of the clitellum) in the tropical endogeic earthworms *Pontoscolex corethrurus* (invasive) and *Balanteodrilus pearsei* (native) after 100 days of culture under the interaction of three diets of different nutritional quality and three moisture content levels in soil with low anthropic impact. Vertical lines represent standard error.

### Cocoon Production

*B. pearsei* and *P. corethrurus* displayed biparental and uniparental sexual reproduction, respectively. On *M. pruriens* treatments (25 %, 33 % and 42 % SM), cocoon production started when *B. pearsei* and *P. corethrurus* reached a mean biomass of 773.5 ± 146.8 mg and 644.7 ± 71.1 mg (average of 25 %, 33 % and 42 % SM), respectively. On maize stubble treatments, it started when *B. pearsei* and *P. corethrurus* reached a mean body weight of 593.0 ± 80.9 mg and 598.5 ± 95.2 mg (average of 25 %, 33 % and 42 % SM), respectively. Cocoon production in *P. corethrurus* was observed in soil (6.7 %), maize stubble (53.3 %), and *M. pruriens* (86.7 %) treatments, but in *B. pearsei* it was observed only in maize stubble (33.3 %) and *M. pruriens* (86.7 %) treatments.

Mean cocoon production was significantly influenced by EO, CF, SM, and the interaction between these three factors (Table 1). After 100 days of culture, peak mean cocoon production in *B. pearsei* and *P. corethrurus* was observed in *M. pruriens* treatments, with 59.7 ± 40.8 and 35.5 ± 21.5 cocoons (average of 25 %, 33 % and 42 % SM treatments), respectively (Fig. 3). When fed maize stubble, *B. pearsei* and *P. corethrhrus* produced 7.9 ± 3.2 and 14.4 ± 9.2 cocoons (average of 25 %, 33 % and 42 % SM treatments), respectivelly. Finally, when fed soil only (33 % SM), *P. corethrurus* (448 mg body weight) produced only two cocoons.

**Fig. 3.**
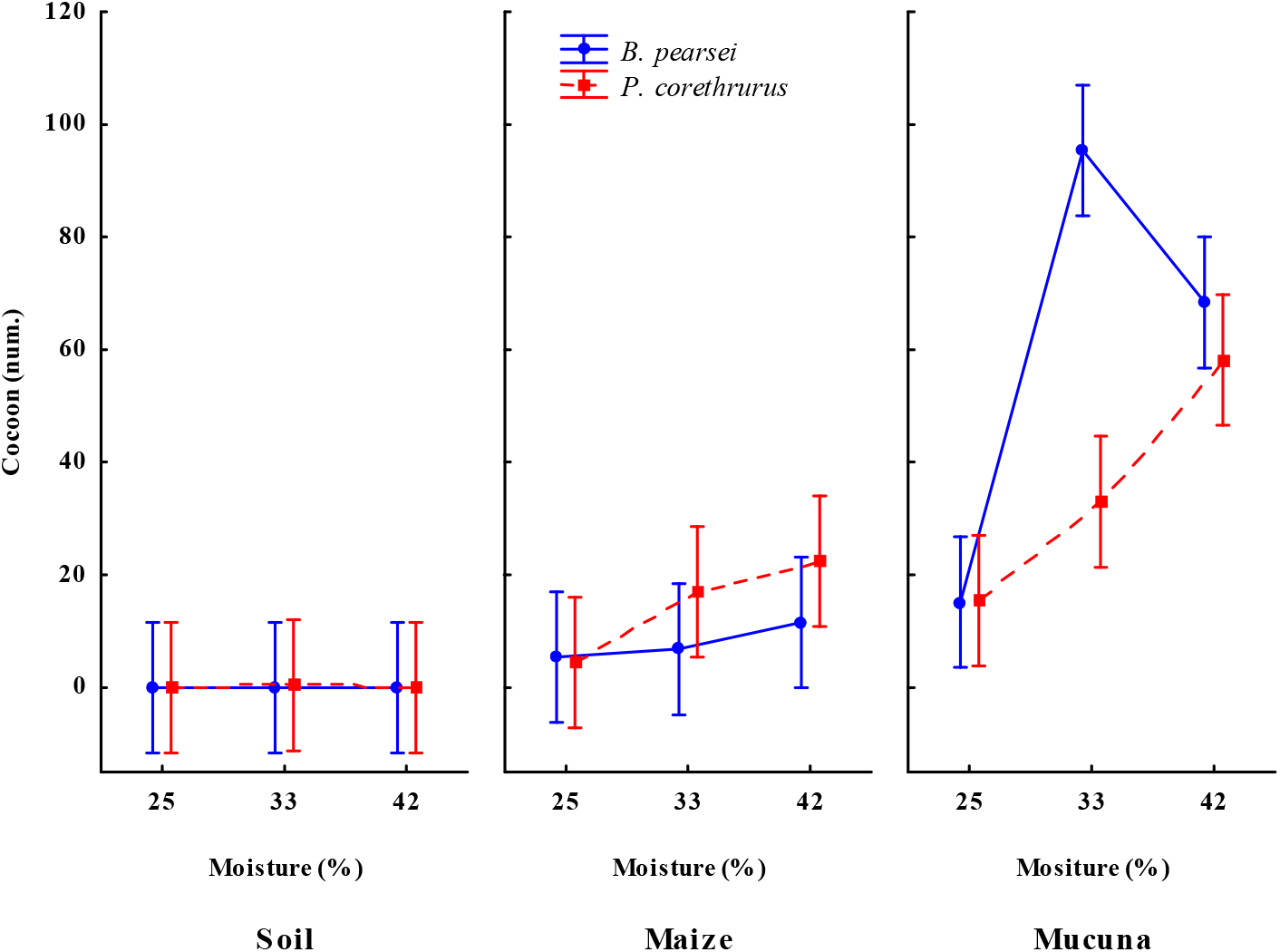
Number of cocoons produced by the tropical endogeic earthworms *Pontoscolex corethrurus* (invasive) and *Balanteodrilus pearsei* (native) after 100 days of culture under the interaction of three diets of different nutritional quality and three moisture content levels in soil with low anthropic impact. Vertical lines represent standard error.

Cocoon biomass varied significantly in response to EO, FQ, SM and the OE x SM and FQ x SM interactions (Table 1). Average cocoon biomass produced by *B. pearsei* and *P. corethrurus* with SM treatments (25 %, 33 % and 42 %) was 10.2 ± 1.4 mg and 27.7 ± 3.7 mg, respectively (Fig. 4).

**Fig. 4.**
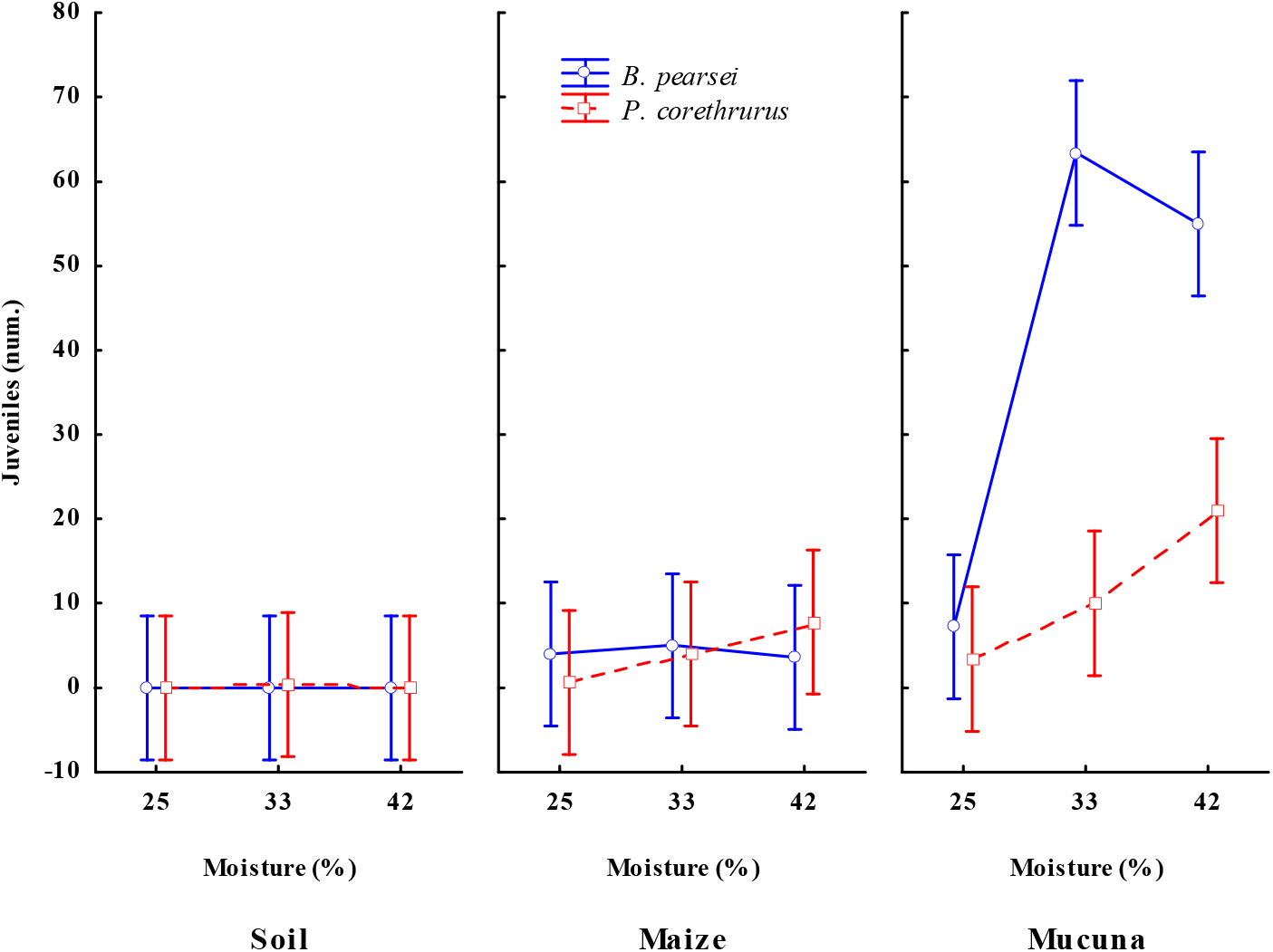
Number of juveniles hatched from cocoons produced by the tropical endogeic earthworms *Pontoscolex corethrurus* (invasive) and *Balanteodrilus pearsei* (native) after 100 days of culture under the interaction of three diets of different nutritional quality and three moisture content levels in soil with low anthropic impact. Vertical lines represent standard error.

### Juvenile Production

The mean cocoon incubation time was similar among treatments (P > 0.05). In general, mean cocoon incubation time was 20.4 ± 5.2 days (*B. perasei*) and 30.3 ± 2.2 days (*P. corethrhrus*), with one individual hatching per cocoon in all cases. Of the total number of cocoons produced by *B. pearsei* and *P. corethrurus* in *M. pruriens* and corn stubble treatments, the average number of hatched juveniles was 64.7 ± 16.6 % and 29.5 ± 7.0 % (average of 25 %, 33 % and 42 % SM treatments) and 59.5 ± 24.7 and 24.0 ± 10.6 (average of 25 %, 33 % and 42 % SM treatments), respectively.

The number of hatched juveniles of *B. pearsei* and *P. corethrurus* varied significantly with OE, CF, SM, and the interaction between these three factors (Table 1). The mean number of hatched juveniles of *B. pearsei* and *P. corethrurus* increased in adults fed *M. pruriens*, as well as with increasing soil moisture (mean 59.7±40.8 and 35.5±21.5 individuals, respectively), and corn stubble (mean 7.9 ± 3.3 and 14.6 ± 9.2 individuals) (Fig. 5). In the *M. pruriens* and corn stubble treatments (Fig. 6), mean biomass of *P. corethrurus* juveniles (21.2 ± 1.0 and 18.6 ± 7.4 mg, respectively) was higher vs. *B. pearsei* juveniles (8.5 ± 0.7 and 8.5 ± 1.3 mg, respectively).

## Discussion

When domesticated, wild populations respond to changing selective pressures, which are reflected in their adaptation to agricultural niches (Boivin et al 2016; Stitzer et al. 2018). From an ecological perspective, the endogeic earthworm *P. corethrurus* resembles non-domesticated species or weeds given its strong profile (invading species) regarding growth rate, fertility, plasticity, interspecific competition, and environmental tolerance (Blakemore 2009; Willcox 2012; Vigueira et al. 2013; Mercuri et al. 2018). In the present study, soil in the habitat for *B. pearsei* was observed to restrain the presence of *P. corethrurus*, a finding consistent with the absence of *P. corethrurus* in parcels where maize- and *M. pruriens* crop rotation is practiced, as well as in pastures and other traditional tropical agroecosystems (Lavelle et al. 1981; Ortiz-Ceballos et al. 2004; Huerta et al. 2006; Marichal et al. 2010; Fragoso et al. 2016; Ortiz-Gamino et al. 2016).

The conversion of the Amazon forest to pastures led the homogenization of bacterial communities in soil, wich differ from those in the adjacent forest, i.e., beta diversity was reduced (Fierer et al. 2013; Rodrigues et al. 2013; Pérez-Jaramillo et al. 2016). The potential resistance of soil (i.e., predators, low species richness, etc.) to earthworms has been documented (Gilot-Villenave 1994; Verstergård et al. 2015; Menezes et al. 2018); for instance, it has been recorded that the endogeic tropical earthworm *Millsonia anomala* from the savannah was unable to prosper in forest soil (Gilot-Villenave 1994). Also, in a Glenrock soil (southern Australia), the survival of the European endogeic earthworm *Aporrectodea trapezoides* was found in association with soil bacteria (Davidson et al 2013; Menezes et al. 2018). Also, the shift in vegetation from grass to woody plants decreaced in the density and biomass of *P. corethrurus* (Sánchez de León and Zou, 2004). Our results showed that the survival of *P. corethrurus* was lower in the environment where *B. pearsei* thrives, maybe due to a negative interaction with a more diverse edaphic microbiome (Gilot-Villenave 1994; Philippot et al. 2013; Menezes et al. 2018).

Symbiosis, defined as the interaction between two different organisms living in close physical association, has been acknowledged as a source of evolutionary innovation that allows hosts to exploit otherwise inaccessible niches (Laland et al. 1999; Lund et al. 2010b; Aira et al. 2018). The hologenome (sum of the genetic information of the host and its microorganisms) theory of evolution is based on four generalization: a) all animals and plants establish symbiosis with microorganims; b) microorganisms can be transmitted between generations with fidelity; c) symbiosis affects the fitness of holobionts in their environment; d) genetic variation in holobionts can be enhanced by incorporating different symbiont populations and can change rapidly under enviromental stress (Zilber-Rosenberg and Rosenberg 2008). Earthworms harbor symbiotic microbiomes that are essential for their life history in the gut, nephridia (excretory organs), and cocoons, but the evolutionary relationship in tropical species such as *P. corethrurus* is poorly understood (Zachmann and Molina 1993; Schramm et al. 2003; Daane et al. 1999; Lund et al. 2010a and 2010b; Aira et al. 2018); for instance, the microbiome is known to improve the nutritional status of low-quality diets (Lund et al. 2010b; Aira et al. 2018). Our findings showed that *P. corethrurus* and *B. pearsei* differ in their diet preference (*M. pruriens*, corn stubble, and control), i.e., the invasive species displayed a faster growth than the native species when nutritional quality improved. This suggests that *P. corethrurus* consumes and degrades a greater variety of organic materials given its greater ability (efficiency), evidenced by: a) producing endogenous cellulases: Ean-Eg, EF-EG2, and GHF9 (Nozaki et al. 2009; Shweta 2012; Ueda et al. 2014; Park et al. 2017); b) its association with the gut microbiota: Protobacteria, Firmicutes, Actinobacteria, Chloroflexi, and Bacteroidete (Thakuria et al. 2010; Liu et al. 2017; Gong et al. 2018; Liu et al. 2018); c) expressing genes (transcriptome) that contribute to the adaptation of its digestive system: salivation, gastric acid, and pancreatic secretion (Liu et al. 2018); d) improving its digestion efficiency according to the type of cecum (Nozaki et al. 2013; Ikeda et al. 2017); e) its association with nephridial Pedobacter bacteria (Davidson and Stahl 2006; Davidson et al. 2013; Menezes et al. 2018).

It is known that in diets of low nutritional quality, mutualistic bacteria (Bacteroidetes, Betaproteobacteria and Alphaproteobacteria) residing in earthworm nephridia (in 19 of 23 species studied) provide vitamins to its host, stimulate earlier sexual maturity, and contribute to pesticide detoxification (Davidson et al. 2013; Lund et al. 2010a, 2010b; Møller et al. 2015; Ponesakki et al. 2017; Aira et al. 2018). The results reported here showed that a lower nutritional quality of the diet (*M. pruriens* > corn stubble > soil) translated into invasive species of smaller size (biomass) that reached sexual maturity earlier than the native earthworm. This suggests that the nephridial symbionts of *P. corethrurus* are generalists, while those of *B. pearsei* are specialists.

Earthworms produce external cocoons that are colonized by bacteria from parents and soil, (vertical and horizontal transmission, respectively; Zachmann and Molina 1993; Aira et al. 2018) and coul be used as biovector for the introduction of benefical bateria (Daane et al. 199). In a new habitat, cocoons of invasive earthworms may be affected by the native microbiota, but they can survive if they carry a parental microbial inoculum. In *Eisenia andrei* and *E. fetida*, 275 and 176 bacteria were observed in their cocoons, respectively (Menezes et al. 2018); however, these were dominated by three vertically transmitted (parental) symbionts: Microbacteriaceae, *Verminephrobacter*, and *Ca. Nephrothrix.* For example, in a sterile environment, cocoons of *M. anomala* failed to develop, suggesting a functional relationship between nephridial bacteria of earthworms and soil microbiota (Gilot-Villenave 1994). Besides, it has been noted that during embryogenesis, the nephridial symbiont *Verminephrobacter* (Betaprototeobacteria) contributes to the biosynthesis of vitamins and the elimination of nitrogen wastes from cocoons, and conserving nitrogen (Schramm et al. 2003; Davidson and Stahl 2006; Lund et al. 2010b; Davidson et al. 2013; Møller et al. 2015). Our findings showed that *P. corethrurus* produced cocoons when fed either of the three diets, while *B. pearsei* fed the control diet (only soil) failed to produce cocoons. In contrast, cocoons of *P. corethrurus* had a low hatching rate (births), which was lower (diet with *M. pruriens*) vs. *B. pearsei*. These results suggest the absence and/or loss of parental symbionts bacteria, i.e., the loss of a parental care strategy to control predators, detoxify nitrogenous wastes, conserving nitrogen, and supply vitamins and essential cofactors (Schramm et al. 2003; Lund et al. 2010b; Davidson and Stahl 2006; Davidson et al. 2013; Møller et al. 2015).

The human-mediated translocation of species dates back to the Late Pleistocene (Denevan 1992; Boivin et al. 2016). Invasive plant species are usually divided in two groups according to time of their residence time: archaeophytes brought in up to 1500 AD, and neophytes brought in after this date (Kowarik 1995). Pyšek et al (2005) observed that archeophyte weeds are common in crops introduced at the beginning of agriculture (cereals), but are poorly represented in crops introduced relatively recently (rape, maize), where neophyte weeds are most numerous. This approach can contribute to elucidate the history of the invasion of *P. corethrurus* in Latin America. In Mexico, it is unknown whether *P. corethrurus* arrived before or after the inmigration of Europeans in 1500 AD. It is documented that an exchange of domesticated plants (manioc, maize, cacao and others), as well as recently introduced plants (livestock, pastures, coffee, sugar cane and others), took place between the Amazon area and Mesoamerica through agricultural packages that may have also included *P. corethrurus* as well as weeds, mice, insects, etc. (Denevan 1992; Boivin et al 2016; Levis et al. 208).

Therefore, the four *P. corethrurus* ecotypes described by Taheri et al. (2018b) are likely the result of the selective forces imposed by cultivation, agricultural practices, and industrial and urban activities. That is, this species has evolved to develop resistance to herbicides, pesticides, heavy metals, and hydrocarbons, among others (Taheri et al. 2018b). The above may explain why the paradigm of Beddard (1900) “ …tropical earthworms invade only tropical regions (*Neotropical*) …” is no longer valid, as *P. corethrurus* has been recorded across the five biogeographic regions: Nearctic (Gates 1954, Gates 1972, Blakemore 2009), Palearctic (Omodeo et al. 2003; Blakemore et al. 2006; Reynolds and Jones 2006, Sherlock and Carpenter 2009, Blakemore 2009, Cunha et al. 2014, Rota and Jong 2015; Taheri et al. 2018), Ethiopian (Plisko 2001), Indian (Tsai et al. 2000; Blakemore 2009; Singh et al. 2018; Subedi et al. 2018), and Australian (Blakemore 2009). The tropical microclimatic model of Lavelle et al. (1987) for the growth and development of *P. corethrurus* (20-30 °C) is also not applicable to all four *P. corethrurus* agrotypes (James et al. 2019), since a revision of databases and the literature revealed that this species has been collected in Neotropical latitudes with temperate climate (10-20 °C; 1200 to 2000 m a.s.l.; Köppen 2011; Peel 2007; Volken and Brönnimann 2011) from México (Fragoso 2018; Juárez-Ramón and Fragoso 2014, Ortiz-Gamino et al. 2016), Lesser Antilles (Hendrix et al. 2006), Colombia (Gutiérrez-Sarmiento and Cardona 2014), Brasil (Müller 1857; Bunch et al. 2011; Ferreira et al. 2018; Steffen et al. 2018), Argentina (Mischis and Righi 1999; Mischis 2007), Madagascar (Chapuis-Lardy et al. 2010; Villenave et al. 2010), South África (Plisko 2001, Janion-Scheepers et al. 2016) e India (Singh et al. 2018; Subedi et al. 2018).

The origin of *P. corethrurus* may be related to anthropogenic soil formation (“terras mulatas” and “terras pretas”) and the domestication of manioc (bitter and sweet) and peach palm staple food that facilitated sedentary lifestyles in the Amazon region (Lodge 1993; Glaser et al. 2000; Arroyo-Kalin 2010; Clement et al. 2015; Watling et al. 2018; Levis et al. 2018), and has evolved to the point that we cannot recognize their wild predecessors, as evidenced by the recent designation of the *P. corethrurus* neotype from an anthropogenic environment (Müller 1857; James et al. 2019), and by the ambiguity used for assigning its place of origin (Righi 1984; Dupont et al. 2012).

Cryptic invasions occur frequently, often go unnoticed, and are hard to recognize. Also, it is known that cryptic species may display markedly different responses to a given stimulus or stressor (Liebeke et al. 2014; Schult et al. 2016; Morais and Reichard 2018). Therefore, it is important to classify the four *P. corethrurus* ecotypes (Taheri et al. 2018a; James et al. 2019) as separate evolutionary entities over the residence time, and determine the preference of each ecotype in terms of soil type, culture, and climate, and response to stimulus or stressors (pesticides, herbicides, heavy metals, etc.), among others. To this end, transcriptomes (Rad-Seq) may be used to elucidate the gene flow across cryptic lineages (Puga-Freitas et al. 2015; Schult et al. 2016; Morais and Reichard 2018). In addition, epigenetics could be used to investigate the functional adaptations of *P. corethrurus* lineages (Kille et al. 2013; Liu et al. 2017; Ponesakki et al. 2017; Fernández-Marchán et al. 2018). Last, potential symbionts may be found for each agrotype, including bacteria, nematodes, enchytraeids and other life forms associated with earthworms (Coates 1990; Fernández-Marchán et al. 2018).

Based on the results reported here, we conclude that the lower survival and hatching rates of *P. corethrurus* cocoons (offspring) are associated with its symbiotic bacteria, coupled with a higher diversity of the edaphic microbiome in the agro-ecological niche of *B. pearsei.* This suggests that *P. corethrurus* is an exotic species that thrives far from its natural status, i.e., has no wild ancestry in the study area. For this reason, the likely symbiotic eco-evolution of *P. corethrurus* with the microbiome in its gut, nephridia and cocoons should be explored as a source of biogeography and phylogenetic information (Lund et al. 2010a; Brussaard et al. 2012; Davidson et al. 2013; Møller et al. 2015; Zwarycz et al. 2015; Schult et al. 2016; Fernández-Marchán et al. 2018), i.e., going back to the roots (Philippot et al. 2013; Pérez-Jaramillo et al. 2016).

## Supporting information

Capullos

