## Supplementary material for "*Pontoscolex corethrurus:* a Homeless Invasive Tropical Earthworm?": Capullos

### The GENMOD Procedure

| Model Information | |
|---|---|
| Data Set | WORK.CAPULLOS_TOTAL |
| Distribution | Zero Inflated Poisson |
| Link Function | Log |
| Dependent Variable | capullo |

| | |
|---|---|
| Number of Observations Read | 90 |
| Number of Observations Used | 90 |

| Class Level Information | | |
|---|---|---|
| Class | Levels | Values |
| lombriz | 2 | Bp Pc |
| alimento | 3 | Maiz Mucuna Suelo |
| humedad | 3 | CC Intermed PMP |

| Criteria For Assessing Goodness Of Fit | | | |
|---|---|---|---|
| Criterion | DF | Value | Value/DF |
| Deviance | | 465.3601 | |
| Scaled Deviance | | 465.3601 | |
| Pearson Chi-Square | 80 | 89.6866 | 1.1211 |
| Scaled Pearson X2 | 80 | 89.6866 | 1.1211 |
| Log Likelihood | | 3682.2687 | |
| Full Log Likelihood | | -232.6800 | |
| AIC (smaller is better) | | 485.3601 | |
| AICC (smaller is better) | | 488.1449 | |
| BIC (smaller is better) | | 510.3582 | |

Algorithm converged.

| Analysis Of Maximum Likelihood Parameter Estimates | | | | | | | | |
|---|---|---|---|---|---|---|---|---|
| Parameter | | DF | Estimate | Standard Error | Wald 95% Confidence Limits | | Wald Chi-Square | Pr > ChiSq |
| Intercept | | 1 | -22.6553 | 0.0558 | -22.7646 | -22.5460 | 165112 | <.0001 |
| lombriz | Bp | 1 | 0.5194 | 0.0563 | 0.4090 | 0.6297 | 85.15 | <.0001 |
| lombriz | Pc | 0 | 0.0000 | 0.0000 | 0.0000 | 0.0000 | . | . |
| alimento | Maiz | 1 | 25.1866 | 0.0882 | 25.0137 | 25.3594 | 81551.0 | <.0001 |
| alimento | Mucuna | 0 | 26.3529 | 0.0000 | 26.3529 | 26.3529 | . | . |
| alimento | Suelo | 0 | 0.0000 | 0.0000 | 0.0000 | 0.0000 | . | . |

### The GENMOD Procedure

| Analysis Of Maximum Likelihood Parameter Estimates | | | | | | | | |
|---|---|---|---|---|---|---|---|---|
| Parameter | | DF | Estimate | Standard Error | Wald 95% Confidence Limits | | Wald Chi-Square | Pr > ChiSq |
| **humedad** | CC | 1 | -1.0841 | 0.0965 | -1.2733 | -0.8950 | 126.16 | <.0001 |
| **humedad** | Intermed | 1 | -0.0046 | 0.0588 | -0.1199 | 0.1107 | 0.01 | 0.9374 |
| **humedad** | PMP | 0 | 0.0000 | 0.0000 | 0.0000 | 0.0000 | . | . |
| **Scale** | | 0 | 1.0000 | 0.0000 | 1.0000 | 1.0000 | | |

**Note:** The scale parameter was held fixed.

| Analysis Of Maximum Likelihood Zero Inflation Parameter Estimates | | | | | | | | |
|---|---|---|---|---|---|---|---|---|
| Parameter | | DF | Estimate | Standard Error | Wald 95% Confidence Limits | | Wald Chi-Square | Pr > ChiSq |
| **Intercept** | | 1 | -0.4498 | 0.5399 | -1.5080 | 0.6084 | 0.69 | 0.4048 |
| **lombriz** | Bp | 1 | 0.4921 | 0.5738 | -0.6325 | 1.6167 | 0.74 | 0.3911 |
| **lombriz** | Pc | 0 | 0.0000 | 0.0000 | 0.0000 | 0.0000 | . | . |
| **humedad** | CC | 1 | -0.0095 | 0.6421 | -1.2680 | 1.2490 | 0.00 | 0.9882 |
| **humedad** | Intermed | 1 | -1.5520 | 0.7759 | -3.0727 | -0.0314 | 4.00 | 0.0455 |
| **humedad** | PMP | 0 | 0.0000 | 0.0000 | 0.0000 | 0.0000 | . | . |
