## Supplemental Data 1 for "*Pontoscolex corethrurus:* a Homeless Invasive Tropical Earthworm?"

### The LOGISTIC Procedure

| Model Information | |
|---|---|
| **Data Set** | WORK.MADUREZ |
| **Response Variable** | y |
| **Number of Response Levels** | 2 |
| **Model** | binary logit |
| **Optimization Technique** | Fisher's scoring |

| | |
|---|---|
| **Number of Observations Read** | 1980 |
| **Number of Observations Used** | 1980 |

| Response Profile | | |
|---|---|---|
| **Ordered Value** | **y** | **Total Frequency** |
| **1** | 1 | 598 |
| **2** | 0 | 1382 |

**Probability modeled is y=1.**

| Class Level Information | | | |
|---|---|---|---|
| **Class** | **Value** | **Design Variables** | |
| **lombriz** | **Bp** | 1 | |
| | **Pc** | 0 | |
| **humedad** | **1** | 1 | 0 |
| | **2** | 0 | 1 |
| | **3** | 0 | 0 |
| **alimento** | **1** | 1 | 0 |
| | **2** | 0 | 1 |
| | **3** | 0 | 0 |

| Model Convergence Status |
|---|
| Quasi-complete separation of data points detected. |

**Warning:** The maximum likelihood estimate may not exist.

**Warning:** The LOGISTIC procedure continues in spite of the above warning. Results shown are based on the last maximum likelihood iteration. Validity of the model fit is questionable.

### The LOGISTIC Procedure

**Warning:** The validity of the model fit is questionable.

| Model Fit Statistics | | |
|---|---|---|
| Criterion | Intercept Only | Intercept and Covariates |
| AIC | 2427.763 | 819.932 |
| SC | 2433.353 | 926.158 |
| -2 Log L | 2425.763 | 781.932 |

| Testing Global Null Hypothesis: BETA=0 | | | |
|---|---|---|---|
| Test | Chi-Square | DF | Pr > ChiSq |
| Likelihood Ratio | 1643.8309 | 18 | <.0001 |
| Score | 1066.2343 | 18 | <.0001 |
| Wald | 326.9750 | 18 | <.0001 |

| Joint Tests | | | |
|---|---|---|---|
| Effect | DF | Wald Chi-Square | Pr > ChiSq |
| lombriz | 1 | 1.1977 | 0.2738 |
| humedad | 2 | 30.6436 | <.0001 |
| alimento | 2 | 13.4857 | 0.0012 |
| tiempo | 1 | 323.0695 | <.0001 |
| lombriz*humedad | 2 | 19.2528 | <.0001 |
| lombriz*alimento | 2 | 1.8620 | 0.3942 |
| humedad*alimento | 4 | 3.0342 | 0.5521 |
| lombri*humeda*alimen | 4 | 15.0633 | 0.0046 |

**Note:** Under full-rank parameterizations, Type 3 effect tests are replaced by joint tests. The joint test for an effect is a test that all the parameters associated with that effect are zero. Such joint tests might not be equivalent to Type 3 effect tests under GLM parameterization.

| Analysis of Maximum Likelihood Estimates | | | | | | | | |
|---|---|---|---|---|---|---|---|---|
| Parameter | | | | DF | Estimate | Standard Error | Wald Chi-Square | Pr > ChiSq |
| Intercept | | | | 1 | -5.2234 | 0.4238 | 151.9441 | <.0001 |
| lombriz | Bp | | | 1 | -0.4808 | 0.4393 | 1.1977 | 0.2738 |
| humedad | 1 | | | 1 | 2.5103 | 0.4597 | 29.8158 | <.0001 |
| humedad | 2 | | | 1 | 0.8631 | 0.4404 | 3.8405 | 0.0500 |
| alimento | 1 | | | 1 | -17.3064 | 168.2 | 0.0106 | 0.9180 |
| alimento | 2 | | | 1 | -1.6501 | 0.4495 | 13.4762 | 0.0002 |
| tiempo | | | | 1 | 0.0949 | 0.00528 | 323.0695 | <.0001 |
| lombriz*humedad | Bp | 1 | | 1 | -0.2032 | 0.6230 | 0.1064 | 0.7443 |
| lombriz*humedad | Bp | 2 | | 1 | 2.3262 | 0.6339 | 13.4674 | 0.0002 |

#### The LOGISTIC Procedure

**Warning:** The validity of the model fit is questionable.

| Analysis of Maximum Likelihood Estimates | | | | | | | | |
|---|---|---|---|---|---|---|---|---|
| Parameter | | | | DF | Estimate | Standard Error | Wald Chi-Square | Pr > ChiSq |
| lombriz*alimento | Bp | 1 | | 1 | 0.4808 | 237.8 | 0.0000 | 0.9984 |
| lombriz*alimento | Bp | 2 | | 1 | -0.8669 | 0.6353 | 1.8620 | 0.1724 |
| humedad*alimento | 1 | 1 | | 1 | -2.5103 | 237.8 | 0.0001 | 0.9916 |
| humedad*alimento | 1 | 2 | | 1 | -0.3806 | 0.6246 | 0.3714 | 0.5423 |
| humedad*alimento | 2 | 1 | | 1 | -0.8631 | 237.8 | 0.0000 | 0.9971 |
| humedad*alimento | 2 | 2 | | 1 | 0.6910 | 0.6230 | 1.2300 | 0.2674 |
| lombri*humeda*alimen | Bp | 1 | 1 | 1 | 0.2032 | 336.3 | 0.0000 | 0.9995 |
| lombri*humeda*alimen | Bp | 1 | 2 | 1 | -0.1852 | 0.8902 | 0.0433 | 0.8352 |
| lombri*humeda*alimen | Bp | 2 | 1 | 1 | -2.3262 | 336.3 | 0.0000 | 0.9945 |
| lombri*humeda*alimen | Bp | 2 | 2 | 1 | -3.1366 | 0.9079 | 11.9344 | 0.0006 |

| Odds Ratio Estimates | | | |
|---|---|---|---|
| Effect | Point Estimate | 95% Wald Confidence Limits | |
| tiempo | 1.100 | 1.088 | 1.111 |

| Association of Predicted Probabilities and Observed Responses | | | |
|---|---|---|---|
| Percent Concordant | 97.1 | Somers' D | 0.943 |
| Percent Discordant | 2.8 | Gamma | 0.944 |
| Percent Tied | 0.1 | Tau-a | 0.398 |
| Pairs | 826436 | c | 0.972 |
